# Expression trajectories in the aging male mouse colon transcriptome

**DOI:** 10.64898/2026.10.01.756139

**Authors:** Maja Olecka, Konstantin Riege, Madlen Haase, Robert Schwarz, Changxin Wang, Lena Best, Otto W Witte, Christoph Kaleta, Christiane Frahm, Steve Hoffmann

## Abstract

Characterizing temporal dynamics during aging is crucial to understanding the course of aging-related events. The transcriptome is extensively remodeled with age, but the temporal trajectories of these changes, particularly through midlife, remain poorly defined in many tissues. Here, we address this gap in the male mouse colon. Using a dataset of 83 RNA-Seq samples from five time points during adult mouse lifespan, we show that colon aging is characterized by multiple gene-expression trajectories unfolding in parallel, with a major shift occurring during a mid- to late-life transition marked by a collapse of the unfolded protein response (UPR). Moreover, we report widespread non-monotonic trajectories, including genes that peak or dip in midlife and would be missed by comparisons limited to young versus old ages. We further describe age-dependent dynamics of transposable element-containing transcripts and a nonlinear course of transcriptomic entropy across the lifespan. Importantly, we identify a small set of genes that track chronological age and use only three of them to construct a compact transcriptomic clock, which we validate using an independent RNA-seq dataset and qPCR data. Together, our results indicate that nonlinear patterns dominate the aging colon transcriptome and underscore the value of high temporal resolution for identifying key transitions during aging.

## Introduction

Aging is a complex biological process characterized by changes at all levels of biological organization. In recent years, higher temporal resolution across a significant portion of the lifespan has enabled more precise tracking of age-related dynamics and has uncovered nonlinear trends in human and animal models (1–4). Nonlinear trajectories may indicate critical periods in the lifespan when molecular changes accelerate, decelerate, or reverse, which are not captured by standard linear models or by experimental designs that compare only two age groups. Characterizing the temporal dynamics of aging is crucial for understanding the underlying mechanisms and for designing more precisely timed, tailored interventions.

We previously reported nonlinear age-related trajectories in the mouse colon DNA methylome (5). In this study, we instead examine the dynamics of the corresponding transcriptomic data in greater detail. The transcriptome, the complete set of RNA molecules expressed in a cell or tissue at a specific time, serves as a dynamic readout of the system’s genetic and epigenetic state. Age-related changes in the transcriptome result from both stochastic and deterministic mechanisms (6) and impact processes such as splicing, transcription elongation rate, and the balance of transcripts with different lengths (7–9). Beyond alterations in gene expression levels - which have traditionally been the primary focus of transcriptomic studies - other features of the aging transcriptome, including increased expression of transposable elements (10) and changes in transcriptomic entropy (11), are also known to be associated with aging phenotypes.

Nonlinear trajectories of gene expression during aging have been reported for several mouse tissues (12). Here, we extend this work to the colon, where age-related changes may influence both local tissue homeostasis and systemic processes (13–15), by analyzing high-quality RNA-seq data from mouse samples collected at five distinct time points across the lifespan, each with substantial biological replicate counts to ensure robust statistical power. Beyond characterizing changes in gene expression, we also examine additional features of the aging transcriptome, including the dynamics of transposable element expression and alterations in transcriptomic entropy. Our analyses reveal marked, age-dependent shifts in the colonic transcriptome throughout the lifespan, impacting a wide array of molecular processes and highlighting changes in canonical aging-related pathways.

## Methods

### 1.1. Datasets

The bulk RNA-seq dataset comprised 83 male C57BL/6/Ukj mouse colon samples from five age groups: 3 months (*n* = 16), 9 months (*n* = 16), 15 months (*n* = 16), 24 months (*n* = 17), and 28 months (*n* = 18). The dataset was generated previously (5) and is available in the Gene Expression Omnibus (GEO) under accession number GSE248002. The C57BL/6J/Ukj substrain lacks two common mutations found in the C57BL/6J strain: the DIP686 mutation in the crumbs family member 1 (*Crb1*) gene, and a mutation in the nicotinamide nucleotide transhydrogenase (*Nnt*) gene. An independent male C57BL/6J mouse colon RNA-seq dataset comprising five age groups (8) was downloaded from GEO under accession number GSE141252 and used for external validation of the minimal transcriptomic clock.

### 1.2. RNA-seq data processing

Previously published RNA-seq data of male mice colon and five age groups have been re-analyzed utilizing an in-house developed NGS data analysis pipeline (github.com/hoffmann-lab/rippchen) to obtain significantly differentially expressed genes. Briefly, according to FastQC v0.11.9 reports, we used Trimmomatic (16) v0.39 (5nt sliding window approach, mean quality cutoff 20) for quality trimming and Cutadapt (17) v2.10 for clipping of Illumina Universal adapter sequences as well as poly-A termini (minimum overlap 5nt). Fragments of ribosomal RNAs were artificially depleted utilizing SortMeRNA (18) v4.3.7 and subsequently aligned to the reference genome GRCm38 (Mm10) with segemehl (19) v0.3.5 (splice-aware mode, 95% accuracy), filtered via SAMtools (20) v1.16.4 for uniqueness and properly aligned mate pairs before subjected to reverse stranded quantification by featureCounts (21) v2.0.6 (minimum overlap 10nt) on Ensembl v102 reference annotation. Differentially expressed genes across all pairs of age groups, together with variance stabilized read counts, were computed using DESeq2 (22) v1.34.0 under consideration of library preparation batches as additional confounding factor in the design formula. As a significance cutoff an FDR <= 0.1 was chosen. 30 RNA-seq samples of publicly available large intestine data (GSE141252, single-end, unstranded) have been processed analogously.

### 1.3. Quantification of TE expression by RNA-Seq

#### 1.3.1. Generation of SalmonTE reference index

Sequences of transposable elements (TEs) were extracted from the reference genome based on the RepeatMasker annotation of *Mus musculus* (mm10, based on Repeat Library 20140131, downloaded in January 2020, https://www.repeatmasker.org/genomes/mm10/RepeatMasker-rm405-db20140131/mm10.fa.align.gz) as described in (23) and combined with the gene annotation of *M. musculus* mm10 (v102 from http://ftp.ensembl.org/pub/release-102/fasta/mus_musculus/cdna/Mus_musculus.GRCm38.cdna.all.fa.gz). The resulting sequence file served as input for salmon index generation (parameter: --type quasi -k 31) (24).

#### 1.3.2. Alignment and expression quantification

Preprocessed reads (see “RNA-seq data processing” section) were mapped to the generated index using SalmonTE (v0.4) (25), with the expression measurement type set to count (parameter: --exprtype=count). The resulting count matrix was filtered to retain TE counts only, and all TEs with less than or equal to ten reads in total across all samples were removed. Counts were rounded, and differential expression analysis was performed using DESeq2 (v1.34.0, default settings) (22) to identify differentially expressed TEs (deTEs). TE instances for which DESeq2 calculated an adjusted p-value were considered expressed in all downstream analyses. Raw counts were independently normalized by calculating transcript per million (TPMs).

#### 1.3.3. Characterization of deTEs

Known transcription start sites (TSS) of genes were extended by 1,000 bp in 5′-direction and labeled as promoter. Transposable elements (TEs) overlapping exons were classified as exonic. Among the remaining TEs, those overlapping promoter regions were classified as promoter-associated, and of the remaining set, those overlapping gene bodies were classified as intronic. TEs not overlapping any of these features were labeled as intergenic. Strand information was ignored for all intersections. TEs were assigned host genes based on the overlap with a gene.

For each deTE cluster and TE order, enrichment was tested using Fisher’s exact test on a 2 × 2 contingency table containing the numbers of TE instances from the TE order of interest and from all other orders, both within the cluster and in the remainder of the genomic TE annotation. Odds ratios, 95% confidence intervals, and two-sided p-values were calculated in R using fisher.test. The correlation between TE and host gene expression was calculated using Spearman’s correlation using cor.test R function.

### 1.4. Clustering

Genes and transposable elements (TEs) identified as differential in any pairwise comparison were clustered based on their expression profiles using clust (26) (v1.12.0). Z-score transformation of TPM values was chosen as the normalization method. Because clust does not force all features into clusters, genes/TEs that were not assigned to any cluster were iteratively resubmitted to the clustering algorithm.

For gene expression, 79% of differentially expressed genes (DEGs) were assigned after 7 iterative rounds, resulting in 14 clusters. For TEs, 4 iterative rounds resulted in clustering of 60% of differential TEs into 8 clusters.

### 1.5. Transcriptomic diversity

Transcriptome diversity was calculated using normalized Shannon entropy from TPM values of expressed genes (TPM > 0.1) as described in (27). Briefly, for every expressed gene *i*, we computed its relative abundance *p_i_* as the probability of observing a transcript from that gene in the library. We then calculated Shannon entropy *H* over the set of *p_i_* values. Because *H* ranges from 0 (all transcripts from one gene) to *log_2_*(*G*) (equal contribution of all *G* expressed genes), we report the transcriptome diversity *H_s_* = *H*/*log_2_*(*G*), which ranges from 0 to 1 per sample.

### 1.6. Ingenuity Pathway Analysis

Ingenuity Pathway Analysis (28) (IPA; QIAGEN; 27.06.2025 release) was used to identify enriched canonical pathways and to predict potential upstream regulators. Expressed genes (mean TPM across all samples > 0) were used as the custom reference set. For potential upstream regulator analysis, only direct relationships were considered. For each pairwise differential expression comparison, IPA used the provided fold changes (log ratios) to compute activation z-scores. Upstream regulators/pathways were considered activated when the activation z-score was ≥ 2 and inhibited when ≤ −2. P-values of overlap were adjusted for multiple testing using the Benjamini–Hochberg method.

### 1.7. *Xbp1* spliced / *Xbp1* unspliced

*Xbp1* isoform usage ratio was measured between *Xbp1u* (NM_013842.3) and Xbp1s (NM_001271730.1), the latter harboring an additional intron of 26nt at transcriptomic position 831, genomic at chr11:5524281-5524307 respectively, by quantifying split reads. The ratio of alignments supporting a splice-junction versus read-through were reported.

### 1.8. Single-cell analysis

The processed single-cell RNA-seq file GSE247719_PanSci_11_Colon_adata.h5ad was downloaded from the GEO (accession GSE247719) (29). Cells were grouped according to the cell-type annotations provided by the original authors. Expression of selected genes was visualized using the DotPlot function in Seurat v5.0.2. For each gene and cell type, dot size represents the percentage of cells with detectable expression, whereas dot color represents the average expression scaled across the plotted cell types. Default DotPlot parameters were used, except for the color gradient.

### 1.9. qPCR

qPCR analysis was applied to measure the expression of *Map10*, *Dmrta2* and *Vsig2*. For this purpose, the RNA of 10 animals per age group from the published study (5) were used. Total RNA was transcribed into cDNA with the LunaScript RT SuperMix Kit as described by the manufacturer (New England Biolabs, Frankfurt am Main, Germany). QPCR was performed with LunaUniversal qPCR MasterMix (New England Biolabs, Frankfurt am Main, Germany) and specific mouse primers (biomers.net GmbH, Ulm, Germany, *Map10*: fw: GAAGTCCTGGGAAGAAGCGG, rv: TGGAGGAGGGCAAAATGTGT; *Dmrta2*: fw: CGAGTCCTTTTCCGGGTCG, rv: TCTTTGTCAGCCTCCGAACC; *Vsig2*: fw: ACCTCTCCGTGACTGACTCA, rv: GGTCTCCTTGGGCTCCTTTT). Amplification was performed using a Rotor-Gene 6000 cycler (Qiagen GmbH) applying the following cycle conditions: 1 min polymerase activation (95 °C); 40 amplification cycles (95 °C for 15 s, 60 °C for 30 s) followed by melting curve ramp from 72 °C to 99 °C rising by 0.5 degree per step.

### 1.10. Minimal transcriptomic clock

From the set of nine genes that were significantly differentially expressed in all pairwise comparisons between adjacent time points, three genes (*Map10*, *Dmrta2*, and *Vsig2*) were selected for clock construction based on their strong temporal separation and low within-time point variance. Samples were randomly divided into training (80%) and testing (20%) sets using age-stratified sampling.

Gene expression values (TPM) were transformed as

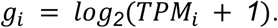

To reduce sensitivity to differences in sequencing depth and global expression scaling between samples, predictor variables were defined as differences in log-transformed expression, corresponding to log-ratios of gene expression:

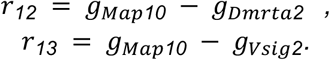

Chronological age was modeled as a continuous response using ordinary least-squares linear regression,

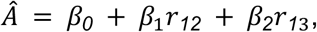

where *Â* is the predicted age and *β0*, *β1*, *β2* are regression coefficients estimated from the training data.

Model performance was estimated by five-fold cross-validation of the training set using the tidymodels framework (v1.3.0). Prediction accuracy was summarized by the coefficient of determination (R^2^) and root mean squared error (RMSE). After cross-validation, the final model was refit using the complete training dataset with the same feature specification and subsequently evaluated on the held-out test set.

To assess model generalizability, the clock was applied without retraining to an independent, publicly available RNA-seq dataset generated from mouse colon tissue (8) (GEO accession GSE141252). Raw sequencing data were processed using the same pipeline as the primary dataset. Because cross-cohort comparisons may be influenced by systematic differences caused for instance by differences in sample preparation or sequencing depth, predicted ages in the external dataset were linearly recalibrated against chronological age before calculation of the final performance metrics. Consequently, this analysis evaluated the model’s ability to capture relative age differences, rather than its ability to provide directly comparable absolute age estimates.

The cross-platform performance of the transcriptomic clock was subsequently evaluated using qPCR data. Predictor variables were calculated from the qPCR cycle-threshold (Ct) values as

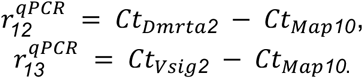

Because qPCR Ct values are inversely proportional to the logarithm of template abundance, these Ct differences provide measures analogous in direction to the log₂-transformed expression ratios used to train the RNA-seq clock. The fitted linear regression model was extracted from the final tidymodels workflow. The qPCR-derived predictors were then supplied directly to this model without refitting the original clock coefficients. As in the external validation analysis, qPCR-derived age estimates were linearly recalibrated against chronological age before calculation of the final performance metrics.

### 1.11. Statistical analysis

All statistical analyses were performed in R (v4.3.1). Group differences were assessed using one-way ANOVA using aov() function, followed by Tukey’s HSD post hoc test using TukeyHSD() for pairwise comparisons. Spearman’s rank correlations were computed using cor.test(), and p-values were adjusted for multiple testing using the Benjamini–Hochberg method. Principal component analysis (PCA) was performed on variance-stabilized counts of 1,000 most variable genes (ranked by variance across samples) to explore global variation among samples. PCA was computed using FactoMineR v2.11 and visualized using factoextra v1.0.7.

### 1.12. Visualization

Plots were visualized using any of the following R packages: ggplot2 v3.5.1, factoextra v1.0.7, eulerr v7.0.2, Seurat v5.0.2.

## Results

### Differential gene expression in aging colon

Investigating nonlinear aging trajectories requires comprehensive data collected at multiple time points across the lifespan. To this end, we analyzed an RNA-seq dataset from male C57BL/6/Ukj mice previously used in a study focused on nonlinear changes in DNA methylation with age (5), which is publicly available in Gene Expression Omnibus (GSE248002). This dataset includes 83 RNA-seq samples spanning five age groups: 3 months (n = 16), 9 months (n = 16), 15 months (n = 16), 24 months (n = 17), and 28 months (n = 18) (**Figure 1a**). Principal component analysis (PCA) confirmed the absence of clear outliers (**Supplemental Figure 1a**).

**Figure 1.**
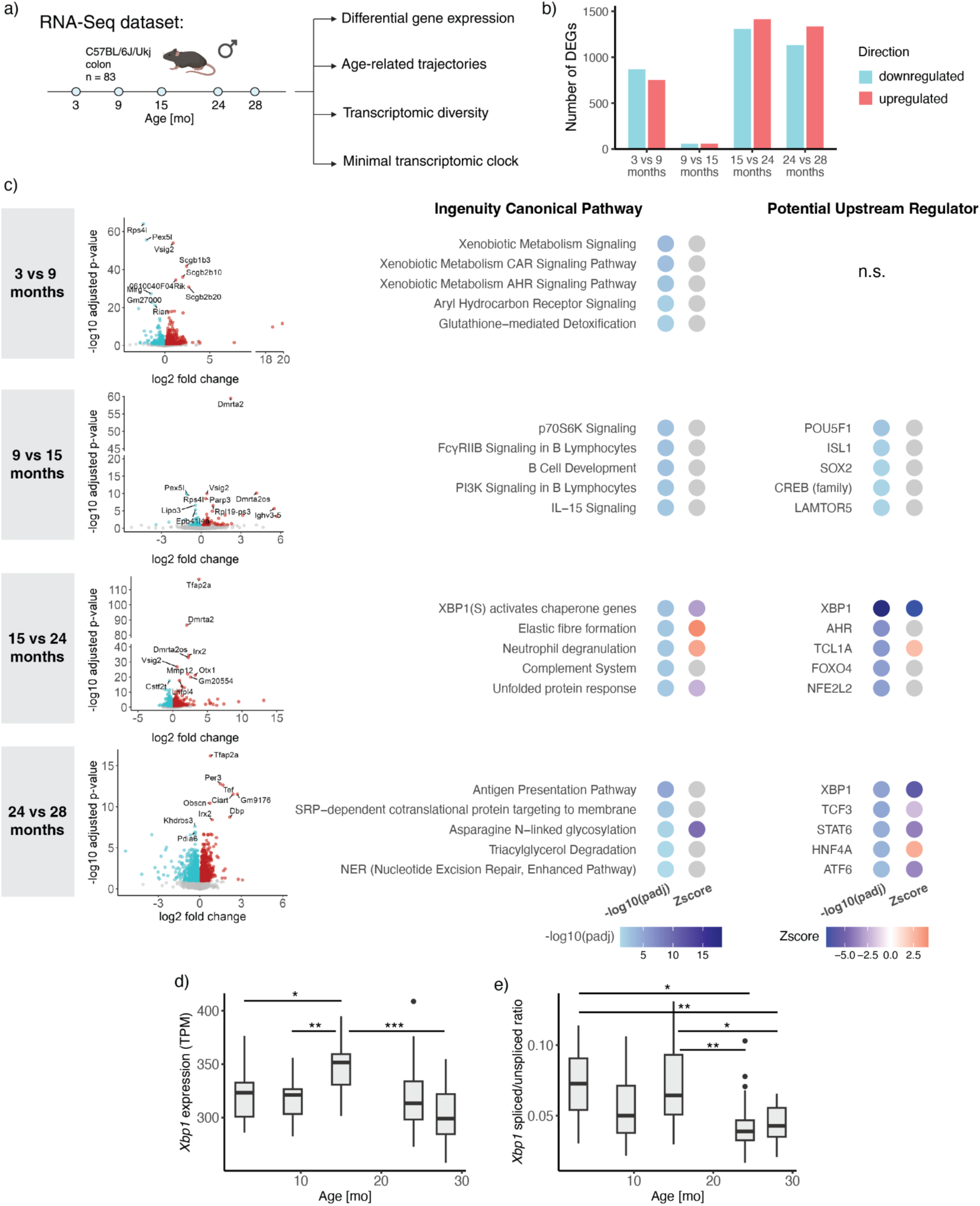
Differential gene expression in the male mouse colon during aging. a) Overview of the dataset and main focus areas of the analysis, created with BioRender. b) Number of differentially expressed genes (DEGs) identified in comparisons between consecutive time points. c) Volcano plots of DEGs and the top predicted upstream regulators and enriched pathways identified in the Ingenuity Pathway Analysis(IPA) analysis. The IPA activation z-score predicts the inferred activation state of a regulator or pathway - positive values indicate predicted activation and negative values indicate predicted inhibition. P-values were adjusted using the Benjamini-Hochberg procedure. n.s., not significant d) *Xbp1* expression during aging. TPM - transcripts per million. e) Ratio of spliced to unspliced *Xbp1* transcripts. In d and e statistical significance was assessed by one-way ANOVA followed by Tukey’s HSD post hoc test. * - P-value < 0.05; ** - P-value < 0.01; *** - P-value < 0.001.

Nonlinear dynamics of the aging transcriptome are already apparent from the number of differentially expressed genes (DEGs). A period of transcriptomic stability between 9 and 15 months, marked by only 116 DEGs, stands in clear contrast to other intervals that show substantially greater numbers of DEGs (**Figure 1b, Supplemental Figure 1b**). Moreover, the majority of DEGs in each comparison, except for the 9 vs. 15-month group, are unique to their respective time points (**Supplemental Figure 1c**), further highlighting the nonlinear nature of gene expression changes in the dataset. Consequently, we detected only 9 genes consistently identified as DEGs across all consecutive comparisons (**Supplemental Figure 1c,d**).

Functionally, the DEGs provide insight into biological processes relevant to distinct life stages (**Figure 1c**). For example, secretoglobulins from the *Scgb1b* and *Scgb2b* gene families, which are upregulated between 3 and 9 months, encode androgen-binding proteins known to influence mate selection in mice (30). *Dmrta2*, upregulated between 9 and 15 months and again between 15 and 24 months, plays a key role in maintaining neural progenitors (31). *Tfap2a* is associated with tumor development (as reviewed in (32)). Additionally, genes involved in circadian regulation, such as *Ciart* and *Per3,* are upregulated between 15 and 24 months and between 24 and 28 months, consistent with previous findings (12,33).

To further characterize gene expression changes across different life stages, we used Ingenuity Pathway Analysis (IPA) to identify pathways and potential upstream regulators associated with differentially regulated genes (**Figure 1c**). The activation z-score was used to predict pathway activation or inhibition based on the directionality of gene expression. Specifically, DEGs between the earliest time points measured in this study (i.e. 3-month vs. 9-month; n = 1,618) are enriched for pathways involved in xenobiotic metabolism and detoxification.

Although only 116 DEGs were identified between 9 and 15 months, we observed clear enrichments. Among these, developmental and stemness regulators, such as POU5F1 (OCT4), SOX2, and ISL1, were identified as potential upstream regulators. Enriched pathways during this period included p70S6K signaling and immune processes.

The comparison between 15 and 24 months, i.e., a mid-to-late-life comparison, yielded the highest number of DEGs (n = 2,722). These genes are enriched in pathways related to ER stress (e.g., XBP1 signaling and the unfolded protein response (UPR)), inflammation (e.g., neutrophil degranulation and the complement system), and extracellular matrix degradation. Notably, FOXO4, a regulator implicated in senescence (34) and colorectal cancer (35), was identified as a potential upstream regulator of the differentially regulated genes during this period. Disrupting the interaction between FOXO4 and p53 has been shown to exert senolytic effects by reducing the viability of senescent cells (36,37).

Between the two oldest time points, i.e., 24 and 28 months, we observed further downregulation of XBP1 target genes and upregulation of antigen-presentation pathways. In addition, pathways related to lipid metabolism, nucleotide excision repair, and cell-cycle control of chromosomal replication were affected during this period. Interestingly, among the strongest signals in our analysis were negative activation z-scores for XBP1- and UPR-associated terms, consistent with a marked inhibition of the unfolded protein response (UPR) between 15 and 24 months and, to a lesser extent, between 24 and 28 months.

The UPR is a central proteostatic mechanism that is triggered by ER stress, i.e., the accumulation of unfolded or misfolded proteins in the endoplasmic reticulum. It comprises three canonical branches mediated by IRE1-XBP1, PERK, and ATF6. Upon activation, these pathways collectively aim to restore proteostasis. To corroborate the predicted downregulation of the IRE1-XBP1 signaling, we quantified the spliced-to-unspliced *Xbp1* transcript ratio (*Xbp1s*/*Xbp1u*), a commonly used proxy for XBP1 pathway activity. This analysis confirmed a decrease in XBP1 activation (**Figure 1d,e**). Because the UPR mitigates ER stress, these findings may appear counterintuitive given that ER stress is generally reported to increase with age (38). However, multiple studies indicate that the capacity to sense proteostatic stress declines with aging and that many UPR components become downregulated as a consequence (reviewed in (39)).

### Age-dependent gene expression trajectories

Pairwise comparisons between age groups offer a clear and interpretable way to assess differences at individual timepoints. However, this approach can overlook broader, nonlinear expression patterns that span the entire aging process. To better characterize these complex dynamics, we clustered genes based on their expression profiles across all time points, grouping those with similar temporal trajectories. Specifically, we included all DEGs identified in any pairwise age comparison (**Supplementary Figure 1b**) and applied the clust algorithm (26) in seven iterative rounds. Ultimately, 11,738 DEGs (79% of all DEGs) were grouped into 14 clusters (**Supplementary Figure 2a, Supplementary Table 1**). For downstream analysis, we focused on clusters containing more than 500 genes, which together comprised 10,021 DEGs (85% of clustered DEGs) (**Figure 2a**).

**Figure 2.**
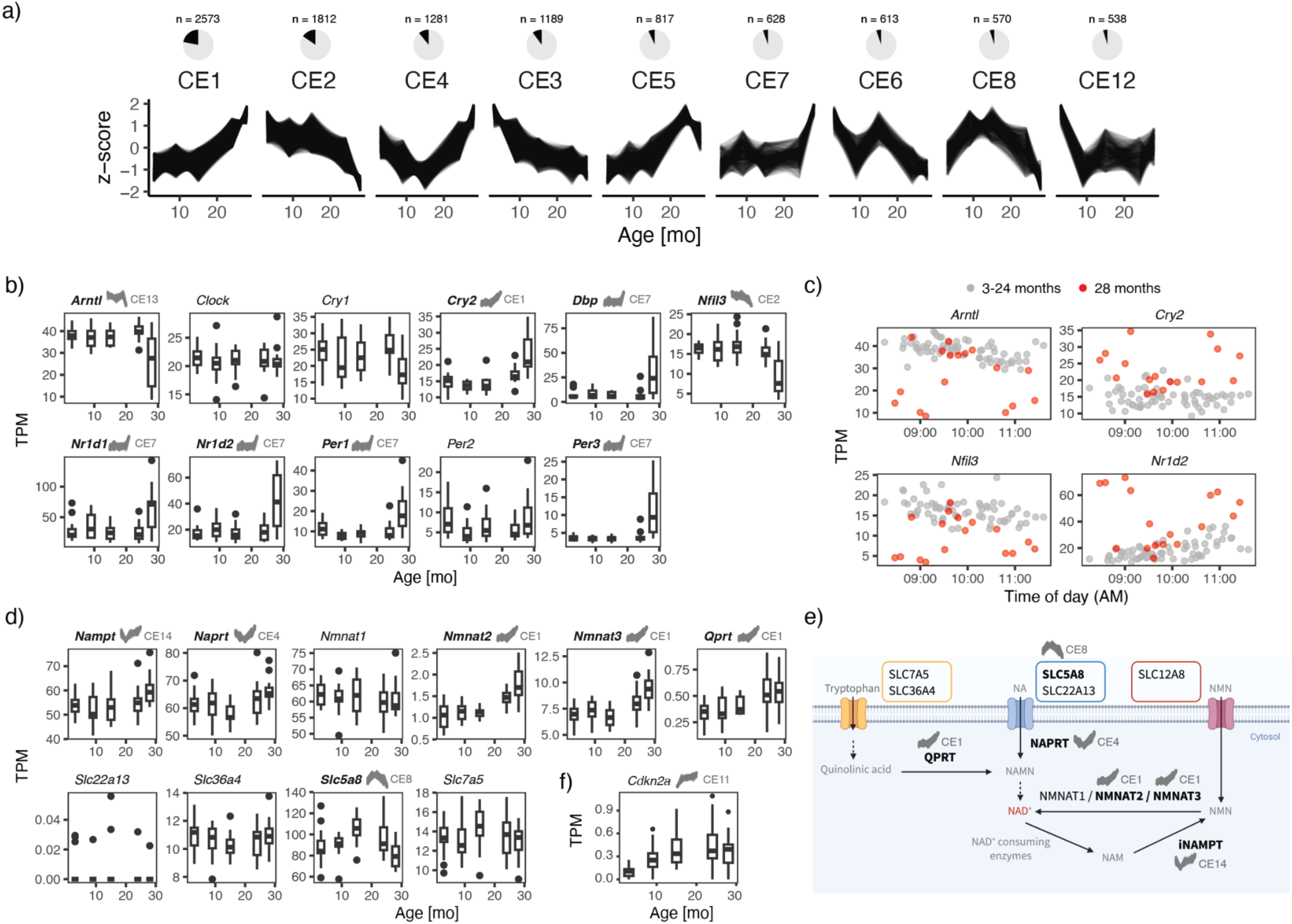
Gene expression trajectories during aging in the mouse colon. a) Largest gene expression clusters (>500 genes per cluster). Pie charts show the proportion of all DEGs assigned to each cluster. All clusters are shown in **Supplementary Figure 2a. b**) Expression of core circadian clock genes during aging and c) examples of dysregulation of day-time dependent gene expression in 28-month-old mice. The x-axis shows the time of sacrifice for each animal. d) Expression of main genes involved in NAD^+^ metabolism. e) Simplified schematic overview of NAD^+^ metabolism based on Li *et al.* (41). DEGs assigned to an expression cluster are shown in bold, with the corresponding cluster name and expression trajectory indicated above each gene, created with BioRender. f) *Cdkn2a* expression during aging. In b, d, f the cluster name and corresponding expression trajectory are shown above the box plot for clustered DEGs.

In contrast to our complementary DNA methylation analysis, where the largest cluster reflects a linear change (5), nonlinear trajectories prevail in gene expression (**Figure 2a, Supplementary Figure 2a**). We detect both monotonic (e.g., CE1, CE3) and non-monotonic (e.g., CE4, CE8) trends. In some cases, the shift in expression is abrupt, occurring early (CE12) or late in life (CE7). The most pronounced shift, represented by the largest clusters CE1 and CE2, occurs between 15 and 24 months, a period we refer to as the mid- to late-life transition. Genes upregulated during this period (cluster CE1) are mainly associated with immune function, consistent with increased inflammation during aging (40), and include glutathione-related genes, which may reflect a compensatory response aimed at maintaining redox balance (**Supplementary Figure 2b**). In contrast, genes downregulated during the mid- to late-life transition (cluster CE2) show higher expression levels than those in cluster CE1 (**Supplementary Figure 2c**) and are associated with mitochondrial function and oxidative phosphorylation, as well as cell-cycle control and ER function (**Supplementary Figure 2b**). Importantly, HNF4A, a key transcription factor in digestive epithelial cells, was identified as a top potential upstream regulator of CE2 genes, suggesting that overall organ functionality may be impaired during this period.

Cluster CE5 follows a trajectory similar to CE1, with an abrupt rise in expression during the mid- to late-life transition (15-24 months). Unlike CE1, however, CE5 shows a subsequent decline in advanced age (24-28 months). Both clusters are enriched for immune-related pathways. CE1 is dominated by terms related to the innate immune system, antigen presentation, and complement, whereas CE5 captures B-cell signaling, stromal remodeling, and elastic fiber formation. The late-life drop of CE5 may reflect B-cell immunosenescence. Alternatively, it could indicate survivor selection, whereby animals with high CE5 gene expression are underrepresented in the oldest cohort.

The nonmonotonic, U-shaped cluster CE4 is enriched in myelination signaling pathway, regulation of the epithelial-mesenchymal transition pathway, and helper T cell activity. Genes with an inverted-U trajectory (CE8) enrich for terms related to granzyme A signaling, pyrimidine de novo synthesis, complex I biogenesis, and triacylglycerol degradation. Lastly, genes showing a zig-zag trajectory (CE6) are enriched, among others, for protein ubiquitination and unfolded protein response pathways.

To illustrate how nonlinear gene expression trajectories can affect aging-related processes, we examined three selected examples with well-established relevance to aging. Two of them point to pathways in which genes follow different trajectories, showing that nonlinear trends within a pathway may not be captured by enrichment analyses restricted to individual expression clusters. The third example focuses on p16, a widely used marker of cellular senescence, to illustrate the age-related trajectory of a canonical aging-associated gene.

Disruption of the circadian clock during aging is common and often precedes the onset or progression of age-related diseases (42). Many circadian clock genes show marked changes in expression across organs with age (33). In our analysis, numerous core circadian clock genes clustered in CE7, showing an abrupt upregulation in the colon between 24 and 28 months (**Figure 2b**). Specifically, *Per1*, *Per3*, *Cry2*, and *Dbp* increased sharply during this interval, whereas genes from the opposing arm of the circadian negative-feedback loop, including the key clock genes *Arntl* (*Bmal1*) and *Nfil3*, were downregulated and clustered in CE13 and CE2, respectively. Importantly, we could not explain this pattern by sampling time. While gene expression in younger animals follows a clear circadian trend, many 28-month-old individuals deviate from this pattern (**Figure 2c**).

Interestingly, we observe that key enzymes involved in nicotinamide adenine dinucleotide (NAD⁺)-associated pathways are upregulated after midlife (**Figure 2d,e**). NAD⁺ is a critical metabolite involved in various cellular processes, including energy metabolism, DNA repair, mitochondrial biogenesis, cellular senescence, and immune cell function. It serves both as a coenzyme in redox reactions and as a cofactor for non-redox NAD⁺-dependent enzymes, such as sirtuins, CD38, and poly(ADP-ribose) polymerases. NAD⁺ levels decline with age in many tissues, a phenomenon that has been associated with age-related diseases (reviewed in (43)). NAD⁺ can be synthesized *de novo* via the kynurenine pathway from tryptophan or through the Preiss-Handler pathway from nicotinic acid, and it can also be regenerated via the nicotinamide (NAM) salvage pathway (44) (**Figure 2e**). Our data indicate that the key salvage pathway enzyme encoding transcript *Nampt* is upregulated after 9 months (CE14), while the downstream enzymes *Nmnat2* and *Nmnat3* increase after 15 months (CE1). Similarly, *Qprt*, the transcript of a kynurenine pathway enzyme, follows a CE1 trajectory. In the Preiss-Handler pathway, *Naprt* displays a U-shaped trajectory (CE4), whereas one of the nicotinic acid transporters shows the opposite, inverted U-shaped trajectory (CE8) (**Figure 2d,e**). Given that age-related NAD⁺ decline has also been reported in the mouse intestine (45), the observed upregulation of synthesis and recycling enzymes may reflect a compensatory mechanism to counteract a NAD⁺ homeostasis decline during aging.

The protein p16 (encoded by *Cdkn2a*) is a cell-cycle regulator that induces senescence by promoting G1 arrest via the retinoblastoma (RB) pathway. Although p16 is widely used as a marker of cellular senescence and aging, suggesting predominant upregulation at advanced ages, our trajectory analysis assigned *Cdkn2a* to CE11, a pattern characterized by a steep increase during early adulthood followed by a more gradual rise after midlife (**Figure 2f**).

Gene expression profiles provide a valuable window into cellular state, although they do not always fully reflect pathway activities and metabolic fluxes. Additional regulatory layers, such as post-translational modifications, protein turnover, or cofactor availability, also shape functional outputs. Our findings, however, highlight the importance of sampling multiple time points in aging studies to better resolve temporal dynamics. When feasible, integrating transcriptomic data with proteomic and metabolomic measurements can further resolve the underlying regulatory mechanisms.

### Age-dependent TE expression trajectories

An important aspect of the aging transcriptome is transposable element (TE) expression. TEs are mobile, repetitive genetic elements that comprise a substantial proportion of the genome. They are usually transcriptionally repressed by epigenetic mechanisms. However, their expression can regulate chromatin accessibility and gene expression and has been shown to be important in development (46). Derepression of TEs has also been implicated in the aging process, in which increased levels of TE transcripts can activate the innate immune system, resulting in chronic inflammation or cause genomic instability through retrotransposition (47). TE-derived transcripts may arise from transcription initiated at a TE’s own promoter (autonomous expression) or from co-transcription with a host gene due to intron retention or transcriptional readthrough (48). Because we used short-read RNA-seq, we were unable to distinguish these sources and therefore, for simplicity, we refer to TE-containing transcripts as “TEs” throughout this study. Nevertheless, both mechanisms have been shown to occur in aging and cellular senescence (49), and analyzing TE-containing transcript dynamics can provide insights into the molecular signatures of transcriptome aging.

TE expression is often quantified at the family level, with reads aggregated across all copies belonging to the same TE family. In contrast, in this study, we adopt a locus-specific strategy (23) and analyze TE expression at the level of individual TE instances. This higher-resolution approach allows us to distinguish differences in expression between copies of the same family. In pairwise comparisons of consecutive time points, we detected the highest number of differentially expressed TEs (deTEs) between 3 and 9 months (**Figure 3a, Supplementary Figure 3a**), and almost all deTEs are unique to their respective pairwise comparisons (**Figure 3b**).

**Figure 3.**
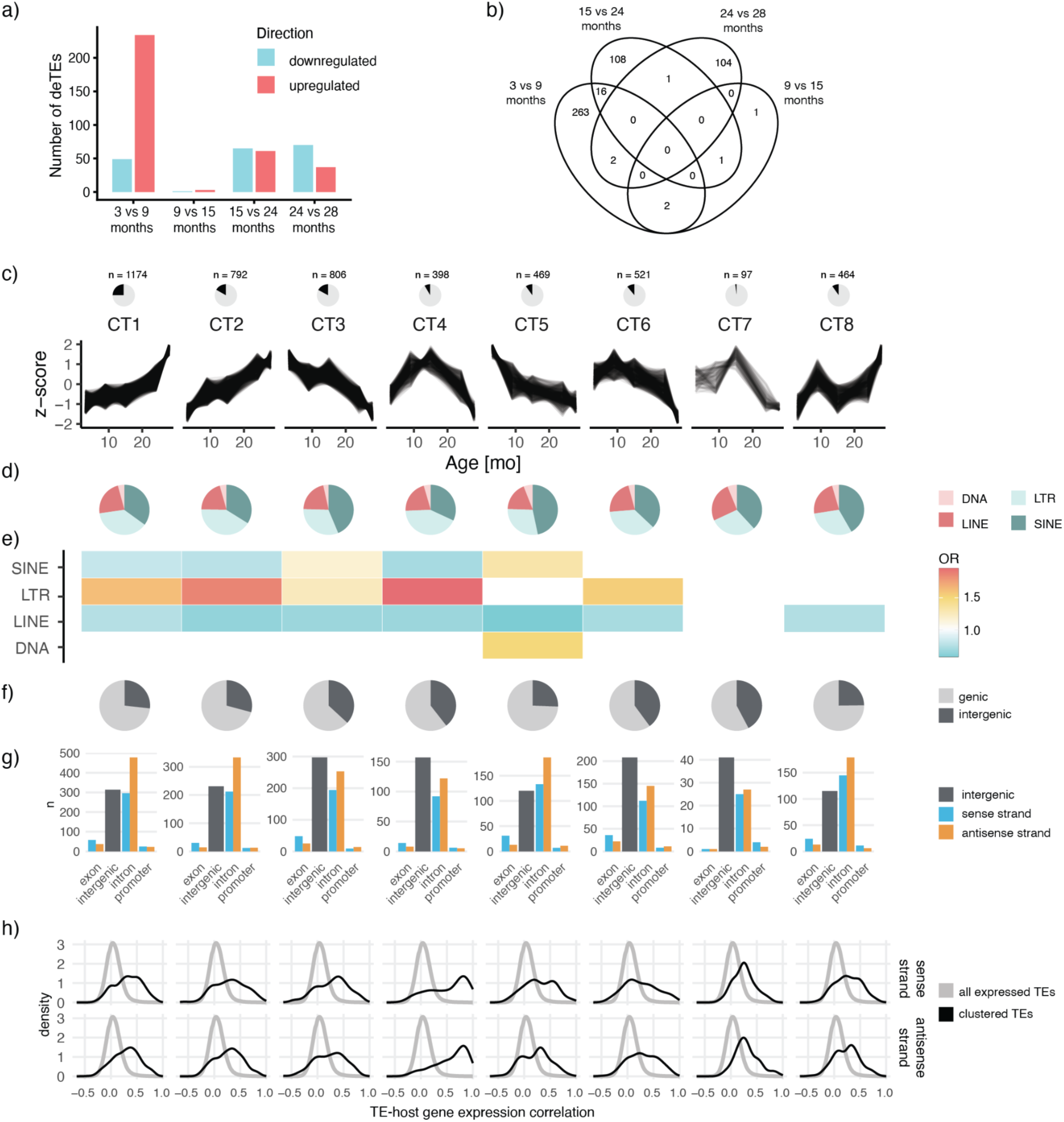
Expression of transposable elements (TEs) during mouse colon aging. a) Number of differentially expressed TEs (deTEs) in comparisons between consecutive time points. b) Overlap of deTEs among comparisons between consecutive time points. c) Expression trajectories of deTEs. Pie charts show the proportion of all deTEs in each cluster. d) Proportion of TEs from different TE orders. e) Enrichment of TEs in different TE orders. OR, odds ratio. f) Proportion of TEs located in genic versus intergenic regions. For comparison, the corresponding proportions for all TEs and all expressed TEs are shown in **Supplementary Figure 3b**. g) Number of TEs in different genomic contexts. For comparison, the corresponding numbers for all TEs and all expressed TEs are shown in **Supplementary Figure 3d**. h) Density plots of the distribution of Spearman’s correlation coefficients between TE and host gene expression. The grey line represents the distribution for all expressed TEs. In g, h sense strand - TE and host gene on the same strand; antisense strand - TE and host gene on opposite strands.

Next, we clustered TE expression trajectories using the same strategy as for genes. After four rounds of clustering, we obtained eight clusters that comprised ∼60% of all deTEs. Although the predominant trend reported in the literature is an age-associated upregulation of TEs, we also observed trajectories reflecting downregulation and non-monotonic patterns (**Figure 3c**). The four largest clusters captured up- and downregulation of TEs that occurred either in an approximately linear manner (CT2, CT3) or in a nonlinear fashion, with transitions in mid- to late-life (CT1, CT6). In addition, we identified a trajectory showing a sharp decline in early adulthood (between 3 and 9 months, CT5), as well as a non-monotonic pattern such as CT4, which displays upregulation in early life followed by downregulation at later age.

Interestingly, distinct TE orders may be differentially engaged at specific phases of life. Specifically, TEs following an early- to mid-life transition pattern (CT5) are enriched for elements from the SINE and DNA transposon orders, in contrast to clusters exhibiting gradual or mid- to late-life transition trends (**Figure 3d,e**). Moreover, the cluster with an inverted U-shaped trajectory (CT4) is strongly enriched for LTR elements.

With the exception of the early- to mid-life transition cluster (CT5), we also observed that downregulated TEs (CT3, CT4, CT6, CT7) tend to have a higher proportion of intergenic TEs (**Figure 3f,g**). This pattern suggests that TE upregulation during aging is often driven by co-expression with host genes, whereas downregulation may more strongly impact TEs transcribed autonomously.

Interestingly, although the U-shaped trajectory cluster CT4 includes a high fraction of intergenic elements, its genic TEs show the highest correlation with host gene expression (**Figure 3h**), and this correlation becomes stronger with age (**Supplementary Figure 3c**). Pathway analysis revealed that these TEs are embedded in genes associated with neuroendocrine and intercellular signaling, involving ERK/MAPK-mediated signal transduction and semaphorin-associated signaling (**Supplementary Figure 4**). Also, the host genes of TEs from the two clusters showing TE upregulation during aging (CT1, CT2) are enriched for age-relevant processes such as immune function, sirtuin signaling, and senescence, and their predicted upstream regulators include chromatin-modifying enzymes (KDM1A, KMT2D), suggesting strong regulation through chromatin modification.

Taken together, our analysis reveals that distinct TE orders and genomic contexts contribute differently to aging-associated TE dynamics, with evidence of both co-expression with host genes and potential changes in autonomous TE activity within specific clusters. To disentangle these mechanisms in more detail, future experiments that directly resolve transcription start sites and autonomous TE transcription, such as CAGE-seq, will be needed.

### Transcriptomic entropy

Moving from locus-level to transcriptome-level analyses, we examined how transcriptomic entropy changes with age. For this analysis, we used Shannon entropy of the gene expression distribution, which quantifies how evenly expression is distributed across genes. High-entropy transcriptomes exhibit relatively uniform expression across genes, whereas low entropy reflects a more skewed distribution, in which a small number of genes dominate total expression while others are weakly expressed or repressed (27).

It was proposed that high transcriptomic entropy is associated with less differentiated, pluripotent cell states, whereas low entropy characterizes highly differentiated, specialized cells (50). Because absolute transcriptomic entropy values are not comparable across samples due to differences in the numbers of expressed genes, we instead use transcriptomic diversity (27), a normalized entropy measure ranging from 0 to 1 (see Methods). Notably, while transcriptomic diversity based on TE expression increases gradually across the lifespan (**Supplementary Figure 5b)**, gene expression–based transcriptomic diversity follows a nonlinear trajectory (**Figure 4a**). It declines between 3 and 9 months, remains relatively stable from 9 to 15 months, and then gradually increases at later ages. This pattern may reflect still ongoing maturation and specialization in early adulthood, followed by age-associated problems with maintenance of cell identity or differentiation capacity in advanced age.

**Figure 4.**
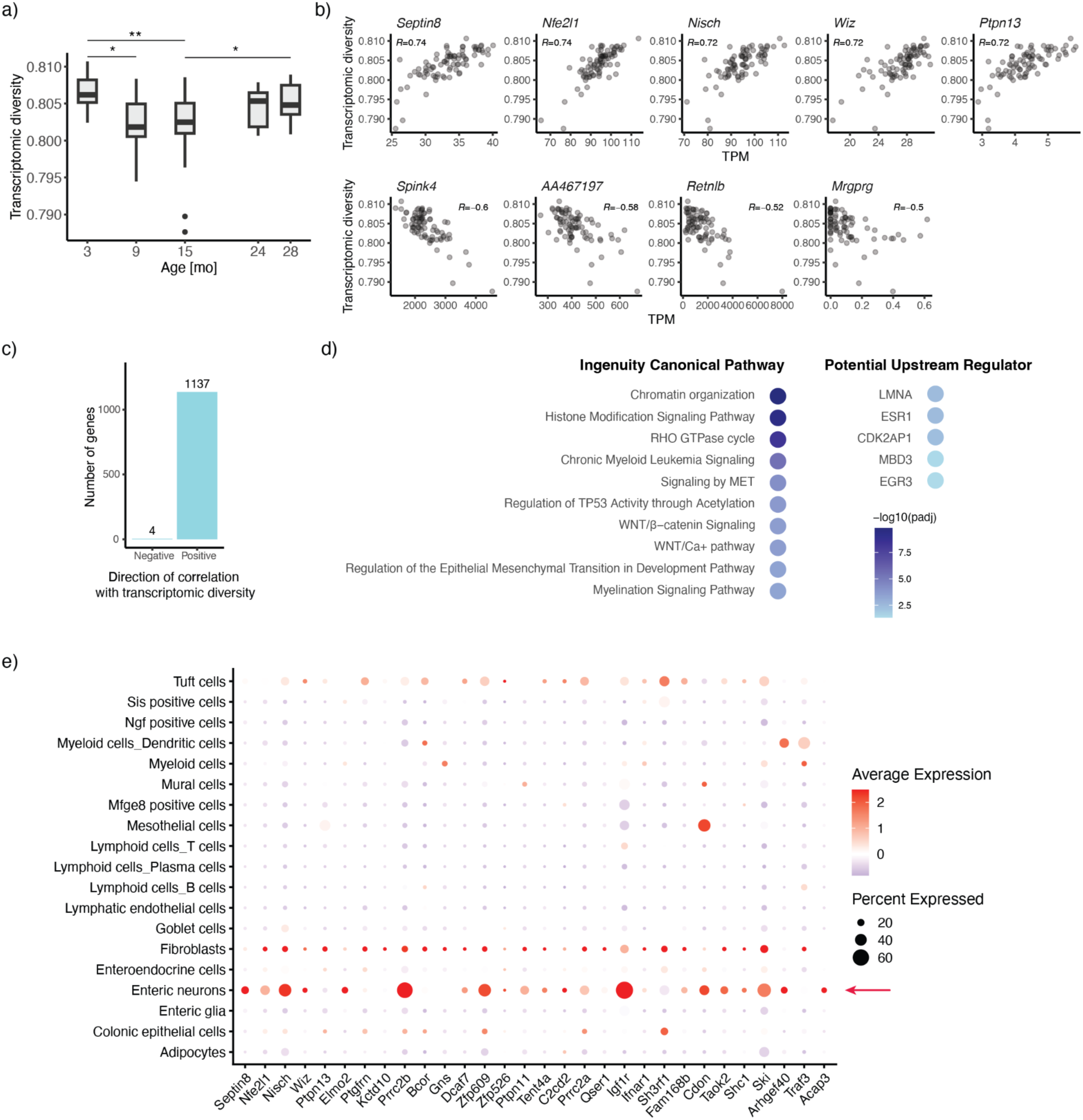
Transcriptomic diversity in the aging mouse colon. a) Nonlinear trajectory of transcriptomic diversity during aging. b) Top genes showing the strongest correlation with transcriptomic diversity (|Spearman’s ρ| ≥ 0.5). c) Number of genes showing positive or negative correlation with transcriptomic diversity. d) Top predicted upstream regulators and enriched pathways identified in the IPA analysis of genes positively correlated with transcriptomic diversity (Spearman’s ρ ≥ 0.5, n = 1,137). P-values were adjusted using the Benjamini-Hochberg procedure. e) Visualization of the expression patterns of the top 30 genes positively correlated with transcriptomic diversity in single-cell RNA-seq data generated by Zhang *et al.* (29). Dot size represents the percentage of cells expressing each gene within the indicated group, and dot color represents the scaled average expression level.

**Figure 5.**
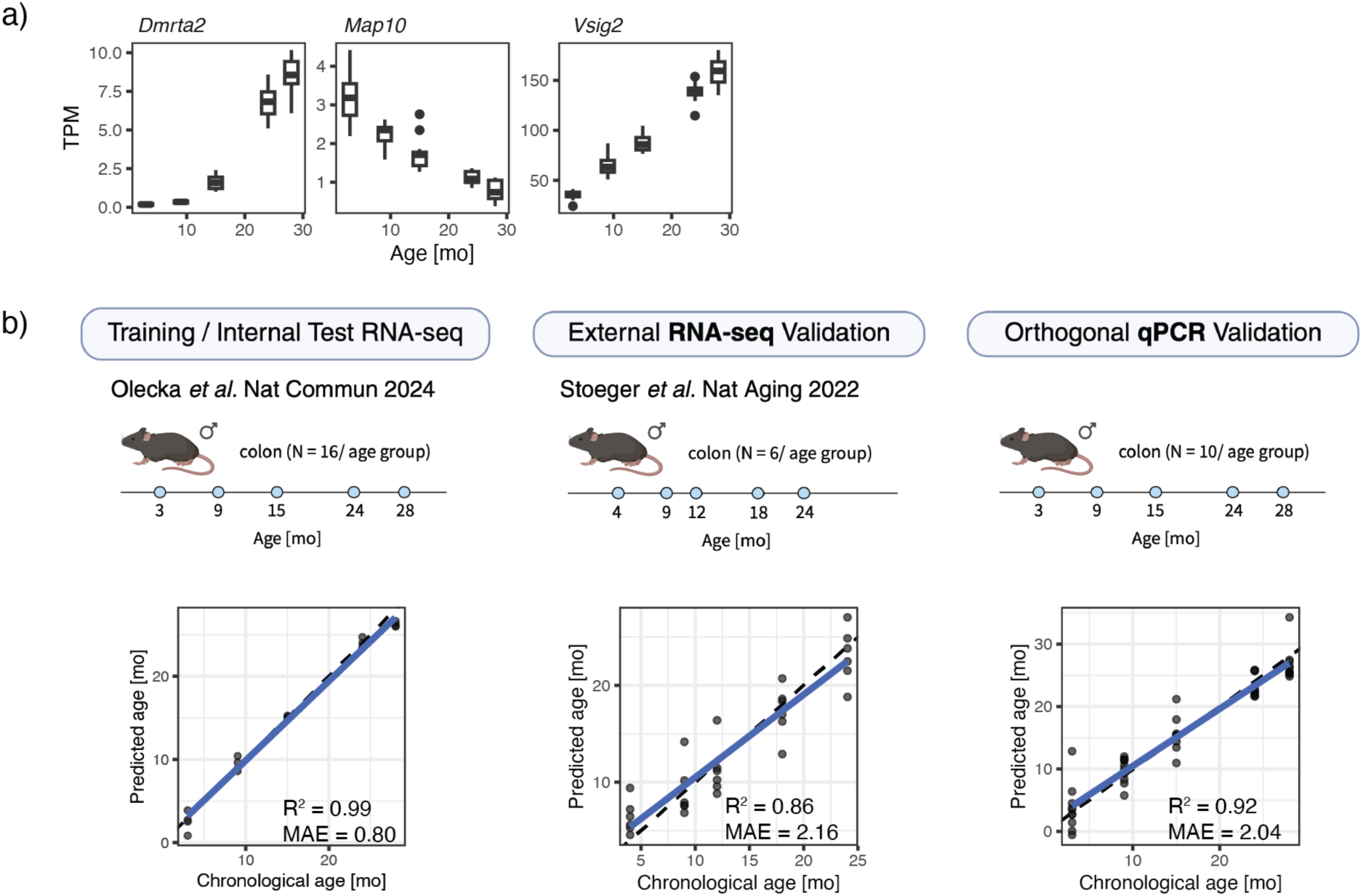
Minimal transcriptomic clock for mouse colon. a) Expression of genes used in the minimal clock during aging. TPM, transcripts per million. b) Datasets used for model development and evaluation (top) and predicted age versus chronological age (bottom). For the external and orthogonal validation datasets, predictions after linear recalibration are shown. Data before recalibration are shown in **Supplementary Figure 6**. The upper panel was created with BioRender. R², squared Pearson correlation between predicted and chronological age; MAE, mean absolute error; dashed line, identity line (y = x); blue line, linear fit.

We then asked which genes correlate with changes in transcriptomic diversity and hypothesized that genes positively correlated with diversity would be enriched for regulators of pluripotency and stem-cell function, whereas negative correlations would mark specialized functions and differentiation programs. Interestingly, we predominantly identified genes showing positive correlation with transcriptomic diversity (Spearman’s ρ > 0.5, n = 1,137), and only 4 genes with negative correlations (Spearman’s ρ < −0.5) (**Figure 4c**). As expected, proteins encoded by the top correlating genes are associated with stem cell function (**Figure 4b**). For instance, WIZ is crucial for maintaining stem cell gene expression programs through structural regulation of DNA loops, together with CTCF and cohesin (51). Moreover, SEPTIN8, a member of the septin protein family, plays important roles in regulating stem-cell quiescence and the balance between symmetric and asymmetric division (52,53).

Furthermore, our analysis revealed that genes positively correlated with transcriptomic diversity are enriched in pathways associated with chromatin organization, histone modifications, and developmental processes, including Wnt signaling and epithelial-mesenchymal transition (EMT) (**Figure 4d**). Strikingly, lamin A (LMNA) was identified as a top potential upstream regulator (**Figure 4d**). LMNA is a principal component of the nuclear lamina and mutations of its gene cause Hutchinson-Gilford progeria syndrome, characterized by accelerated aging (54). These findings are consistent with previous studies suggesting that age-related transcriptomic changes are largely driven by disruption of chromatin architecture (55,56).

We further asked whether genes showing a positive correlation with transcriptomic diversity are cell-type specific or ubiquitously expressed. To address this question, we used single-cell RNA-seq data generated by Zhang *et al.* (29). Visualization of the expression patterns of the top 30 genes positively correlated with transcriptomic diversity suggested that these genes are preferentially expressed in enteric neurons relative to other cell populations, with generally higher average expression levels and detection rates in the enteric neuron cluster (**Figure 4e)**. In contrast, the top negatively correlated genes did not reveal any obvious cell-type-specific expression patterns (**Supplementary Figure 5a).**

Nevertheless, the few genes showing negative correlation with transcriptomic diversity (**Figure 4b**) are associated with intestinal health and epithelial function. *Spink4*, highly expressed in goblet cells, is dysregulated in colitis, and exogenous protein administration alleviates colitis-associated inflammation (57), whereas low *Spink4* levels are linked to poor survival in colorectal cancer (CRC) (58). *Aa467197* contributes to mucosal healing (59), and downregulation of *Retnlb* is associated with reduced survival in CRC (60).

Overall, these results link higher entropy in the colon transcriptome to chromatin remodeling and stem cell programs. In contrast, lower entropy is associated with epithelial identity and intestinal barrier function, suggesting that transcriptomic entropy reflects tissue integrity and differentiation state. Consequently, the higher entropy in aged samples could indicate loss of tissue identity.

### Minimal transcriptomic clock

To extract temporal information from the aging transcriptome, we focused on genes that consistently change across the lifespan. Only nine genes were differentially expressed across all pairwise comparisons between consecutive time points (**Supplementary Figure 1c,d**). Despite complex, nonlinear transcriptional dynamics, these genes showed robust between-time-point differences and low within-time-point variance, indicating strong age dependence and suggesting that a small subset of transcripts encodes temporal information.

Most transcriptomic aging clocks rely on large gene sets and complex machine learning models (61–64). Here, we tested whether chronological age can be inferred using a minimal model based on a small set of informative markers. We selected three genes (*Dmrta2*, *Map10*, *Vsig2*) with strong age association and low intra-time-point variability and trained a linear regression model on log₂-transformed expression ratios (**Figure 7a**). This transformation reduces sensitivity to cross-dataset scaling effects and facilitates translation to qPCR-based assays. The model was trained using five-fold cross-validation on 80% of the cohort and subsequently evaluated on the held-out 20% internal test set, where it predicted chronological age with high accuracy (R² = 0.99, MAE = 0.80 months) (**Figure 7b**).

To test generalizability, we applied the same model to an independent, publicly available RNA-seq dataset from mouse colon generated in a different laboratory (8). In this external cohort, the predictions were strongly correlated with chronological age. However, they were subject to a large bias due to study-specific experimental differences (R² = 0.86, MAE = 10.71 months) (**Supplementary Figure 6**). After recalibration, the model predicted chronological age in the external cohort with good accuracy (R² = 0.86, MAE = 2.16 months) (**Figure 7b**).

To evaluate whether the compact mouse colon aging clock could be transferred to a targeted, low-throughput assay, we applied the model to qPCR-derived Ct values and evaluated its prediction performance. For this analysis, colon RNA samples from mice from the original cohort were used, with 10 animals per age group. Because the original clock was trained on features defined as log₂-transformed TPM ratios, qPCR Ct differences provide an analogous measure of relative gene expression. Similar to the external RNA-seq validation, the qPCR-based model showed an accurate absolute age prediction after recalibration (R² = 0.92, MAE = 2.04 months) (**Figure 7b, Supplementary Figure 6**). These results demonstrate that the compact clock can be transferred to an orthogonal qPCR platform and support its potential use as a targeted assay for assessing colon aging.

Together, while this minimal clock is tissue-specific and was developed in male mice, its simplicity and cross-cohort robustness show that chronological age in the mouse colon can be accurately inferred from the expression of only three genes.

## Discussion

Resolving the temporal dynamics of aging is essential for understanding how molecular and cellular changes accumulate over the lifespan (3). By characterizing the mouse colon transcriptome across lifespan, we show that aging is driven by nonlinear expression trajectories rather than a simple monotonic drift toward an aged state. Importantly, these changes are not restricted to late life. In particular, midlife, a stage often underrepresented in aging studies, exhibits pronounced transcriptional dynamics, with subsets of genes reaching peak or minimal expression. Moreover, the transition from early adulthood to midlife (3 to 9 months) is characterized by multiple changes in gene expression and entropy, indicating that the colonic transcriptome undergoes major remodeling during this period. Notably, the two largest gene clusters are associated with the mid- to late-life transition, suggesting that the most extensive transcriptional reorganization occurs during this window and may define a critical period in colon aging.

Notably, compared with our previous DNA methylation analysis of the same samples (5), nonlinear age-associated patterns were more pronounced in the transcriptome. While the largest cluster in the methylation analysis showed an approximately linear increase with age, the gene expression analysis was dominated by nonlinear trajectories. This likely reflects greater short-term plasticity of the transcriptome and the contribution of regulatory mechanisms beyond CpG methylation. In contrast, age-related methylation changes may capture a smoother, more cumulative component of aging.

The causes of nonlinear aging trajectories remain poorly understood. One plausible mechanism is the age-dependent failure of buffering systems that normally maintain homeostasis under persistent stress (3). Molecular buffers, including proteostasis networks, DNA repair pathways, and defenses against oxidative stress and other sources of damage, allow tissues to withstand damage without immediate phenotypic change. Nonlinearity may emerge when buffering capacities approach saturation: once a critical threshold is crossed, additional damage can no longer be compensated, resulting in a step-like shift in phenotype.

However, rather than a simple depletion of reserve capacity, our data suggest a collapse of the proteostatic buffer itself. Between 15-24 months and 24-28 months, we observe a marked downregulation of XBP1 signaling, despite the expectation that age-related increases in ER stress would activate the UPR. This aligns with prior evidence that UPR sensing mechanisms decline with age and that restoring their activity exerts rejuvenating effects (65–67).

It is important to note that the interpretation of changes occurring between the latest time points analyzed is not straightforward. Gene expression differences observed between 24 and 28 months may not exclusively reflect biological changes associated with aging itself. At these advanced ages, selective mortality may alter the composition of the surviving population, such that mice reaching 28 months represent a more resilient subset of the original cohort. Consequently, some late-life expression patterns may reflect survivorship bias, including expression features associated with increased longevity, rather than changes that occur uniformly during aging.

The application of entropy concepts to aging is not new (68). A prevailing view is that entropy increases with age, yet relatively few studies have tested this hypothesis directly. For instance, it has been reported that organizational tissue structure shows increased entropy with age (69), and epigenetic entropy has likewise been reported to increase in aging tissues (70–72). Consistent with these observations, thermodynamic modeling based on metabolic parameters suggests that calorie restriction reduces daily entropy generation (73). However, dos Santos *et al.* found in human datasets that the dynamics of transcriptional entropy are rather tissue-specific, and that increased transcriptomic entropy has been associated with cancer and age-related disease (11). Our colon data align with the hypothesis that transcriptomic entropy increases with aging. We also show that genes whose expression positively correlates with entropy in the colon play important roles in stem cell biology, suggesting that age-related entropy changes may shift the potential to alter differentiation or maturation states.

Furthermore, we observe that genes positively correlated with transcriptomic diversity are preferentially expressed in enteric neurons, consistent with emerging evidence that intestinal aging involves altered autonomic regulation of the gut. Recent work has implicated intestinal interoceptive dysfunction and impaired vagal afferent signaling in age-associated cognitive decline (74), and broader age-related autonomic imbalance has also been proposed (75). Although our bulk transcriptomic data do not directly measure vagal activity, they are consistent with the possibility that increased transcriptomic diversity reflects disrupted intestinal neural homeostasis.

In contrast, genes negatively correlated with transcriptomic entropy, i.e., genes whose expression decreases as entropy increases during aging are enriched for functions important to colonic physiology, and their down-regulation has been linked to colonic disease. Notably, the relatively small number of genes exhibiting this inverse relationship suggests that the primary constraints maintaining low transcriptomic entropy may operate predominantly at the level of chromatin regulation and epigenetic control.

The small set of genes identified as differentially expressed in all pairwise comparisons revealed transcripts that reliably track temporal progression. Using only three of these genes, we constructed a minimal model that predicts relative age in an independent dataset with high accuracy and, after recalibration, also estimates chronological age. Notably, two of the genes included in the model have previously been implicated in aging-related processes. Specifically, *Map10* has been reported in several murine epigenetic clocks (76–78) and exhibits clear age-associated epigenetic drift (79). This suggests that the locus may capture temporal information at both epigenetic and transcriptomic regulatory levels. MAP10 plays a role in regulating cell division and microtubule stability, raising the possibility that age-associated changes in its expression may reflect processes linked to proliferative history during aging, analogous to the mitotic clock concept proposed for other aging biomarkers. Age-associated changes, although in the opposite direction to changes observed in the colon, have also been reported for *Vsig2*, with protein levels increasing in the plasma of aged mice (80). The overlap between transcriptomic, epigenetic, and proteomic age-associated signals suggests that these loci may reflect fundamental biological processes underlying organismal aging. Taken together, our results indicate that temporal information in the aging transcriptome may be derived from a small subset of highly informative genes, enabling accurate age estimation even with minimal models. Notably, the model also performed well in an independent dataset and qPCR-derived data, which is particularly encouraging given that transcriptomic clocks are generally considered less reproducible across datasets than epigenetic clocks (81). It remains to be determined whether similarly compact signatures can be identified in other tissues and in more heterogeneous settings such as human cohorts.

Taken together, our work supports the view that aging is best conceptualized as a dynamic phenomenon in which multiple molecular processes follow distinct, often nonlinear trajectories. In the aging colon, these trajectories encompass coordinated changes in gene expression, TE activity, and transcriptional entropy, rather than a simple, linear drift away from a young state. Future studies with refined temporal resolution and longitudinal sampling will be essential to resolve these trajectories in greater detail and to identify the key drivers of the nonlinear shifts we observe.

## Author Contributions

M.O. and S.H. designed the study. M.O., K.R., R.S., C.W., and L.B. performed the data analysis. M.H. performed the wet-lab experiments. M.O. wrote the manuscript with critical input from all authors. S.H., C.F., and C.K. acquired funding. S.H., C.F., O.W., and C.K. supervised the study.

## Acknowledgments

The FLI is a member of the Leibniz Association and is financially supported by the Federal Government of Germany and the State of Thuringia.

## Data Availability Statement

The main dataset was generated previously (5) and is available in the Gene Expression Omnibus (GEO) under accession number GSE248002. An independent RNA-seq dataset (8) used for validation is available in the GEO under accession number GSE141252.

