## Supplementary Figures for "Expression trajectories in the aging male mouse colon transcriptome"

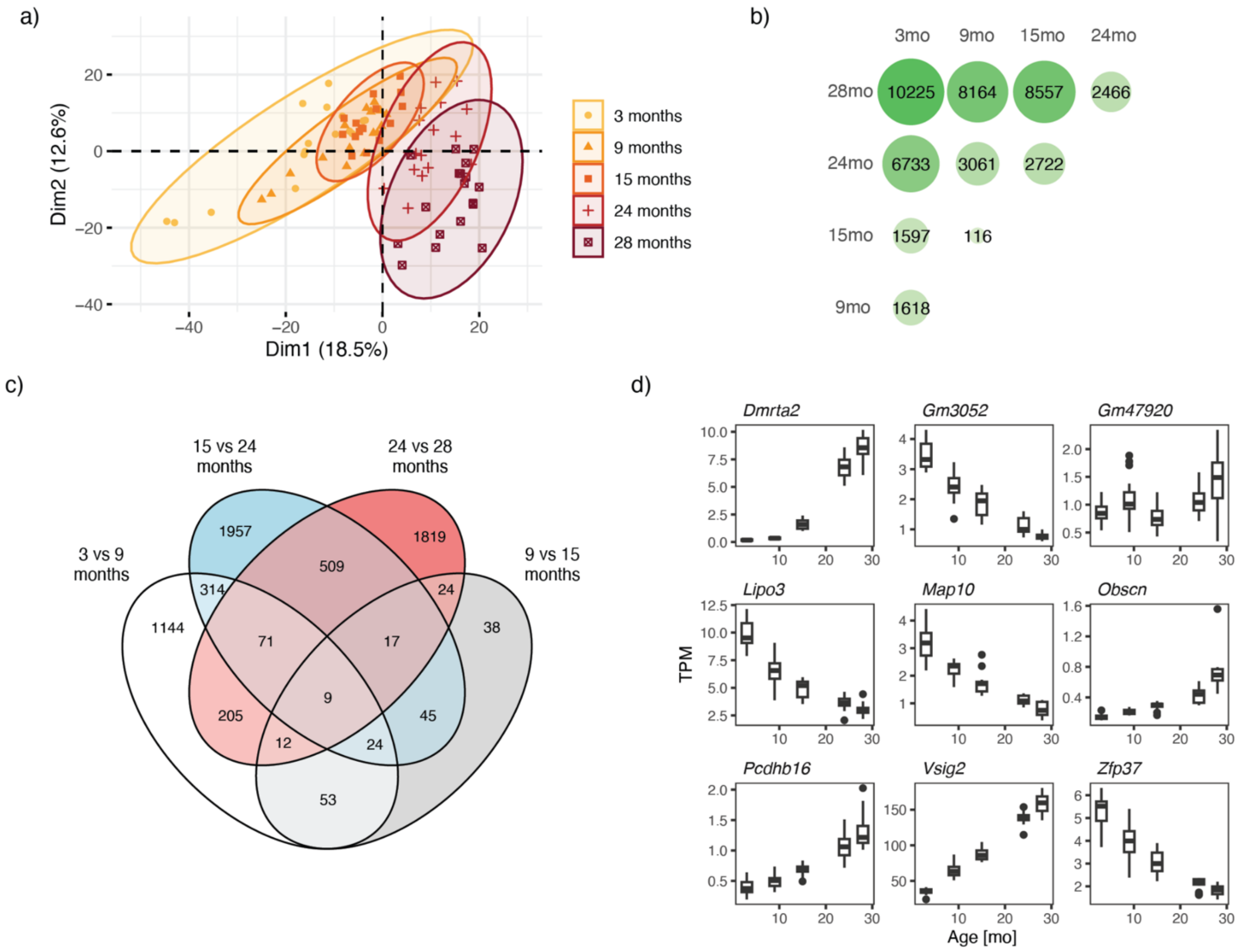


**Supplementary Figure 1.** a) Principal component analysis (PCA) based on 1,000 most variable genes. b) Number of differentially expressed genes (DEGs) identified in all pairwise comparisons. c) Overlap of DEGs between pairwise comparisons of consecutive time points. d) Expression levels of nine genes differentially expressed across all consecutive time point comparisons. TPM, Transcripts Per Million.

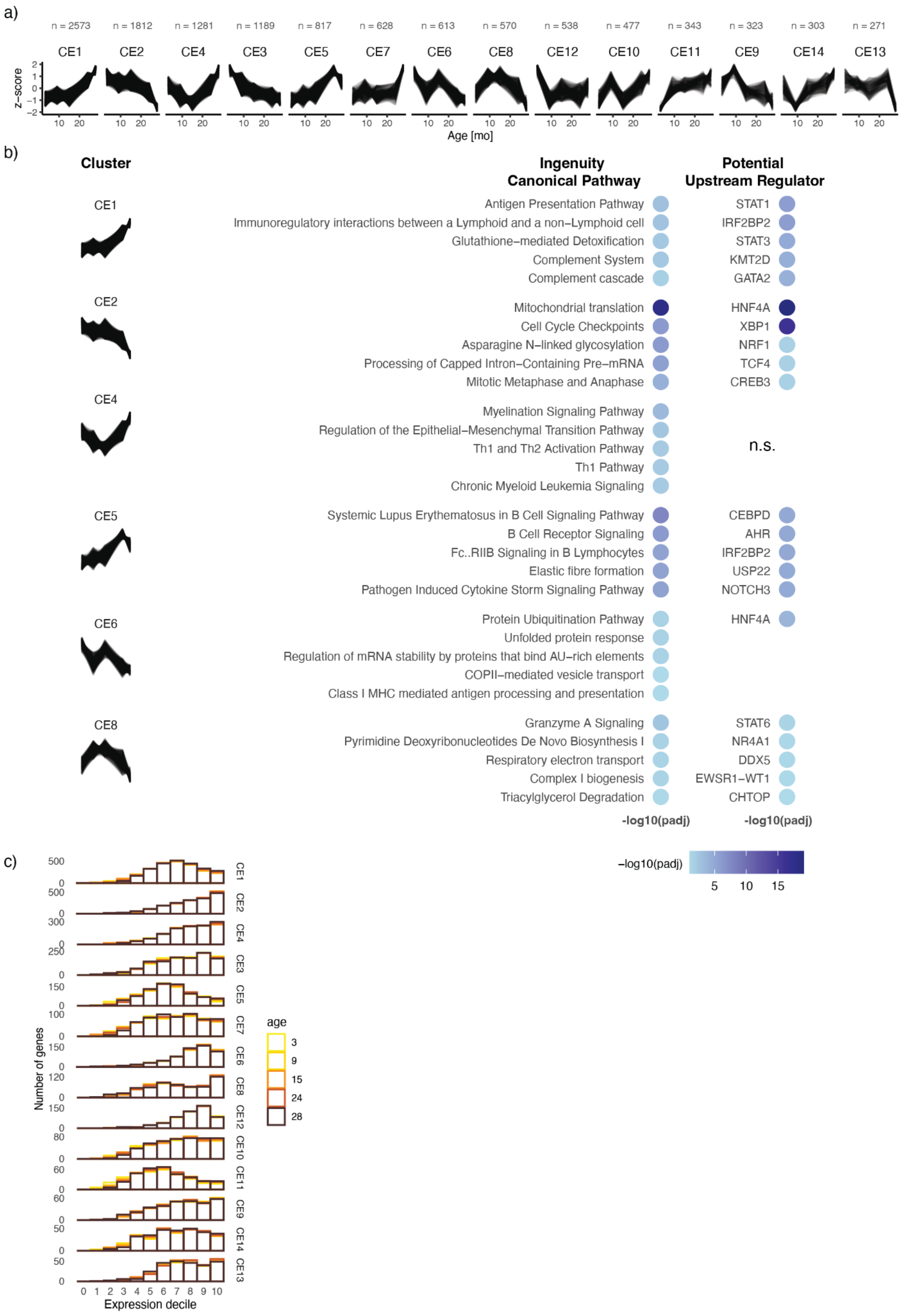


**Supplementary Figure 2.** a) Gene expression clusters ranked by the number of assigned genes. b) Top canonical pathways and predicted upstream regulators identified by Ingenuity Pathway Analysis for genes in the clusters described in the main text. Only results significant after Benjamini-Hochberg correction are shown. n.s., not significant. c) Distribution of gene expression levels across clusters. Expressed genes were divided into deciles, with the lowest expression in decile 1 and the highest in decile 10. Inactive genes were assigned to decile 0.

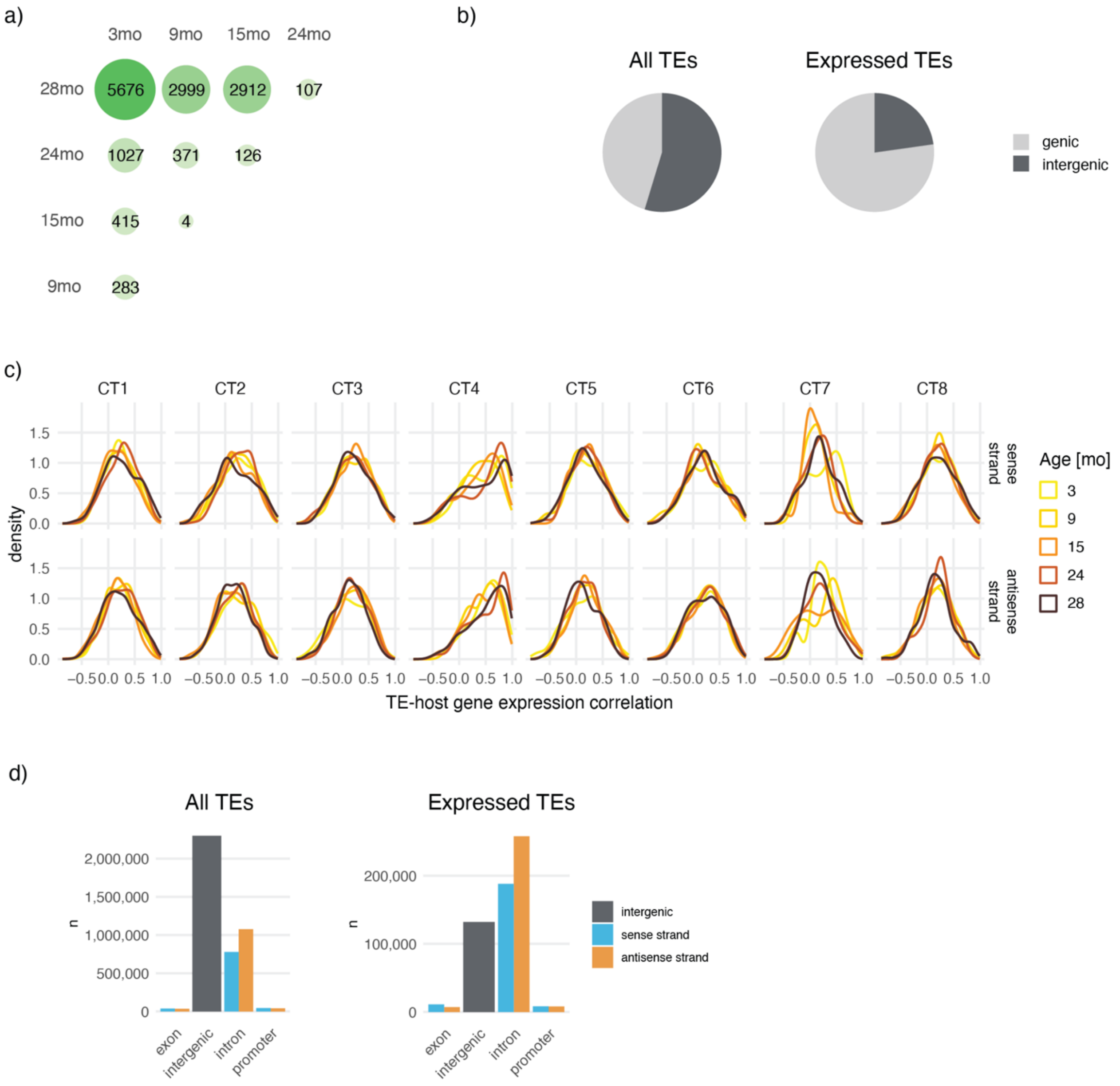


**Supplementary Figure 3**. a) Number of differentially expressed transposable elements (deTEs) identified across all pairwise comparisons. b) Proportions of all TEs and expressed TEs located in genic and intergenic regions. c) Density distributions of Spearman's correlation coefficients between TE and host-gene expression, shown separately for each age group. d) Numbers of all TEs and expressed TEs in different genomic contexts. In c, d sense strand - TE and host gene on the same strand; antisense strand - TE and host gene on opposite strands.

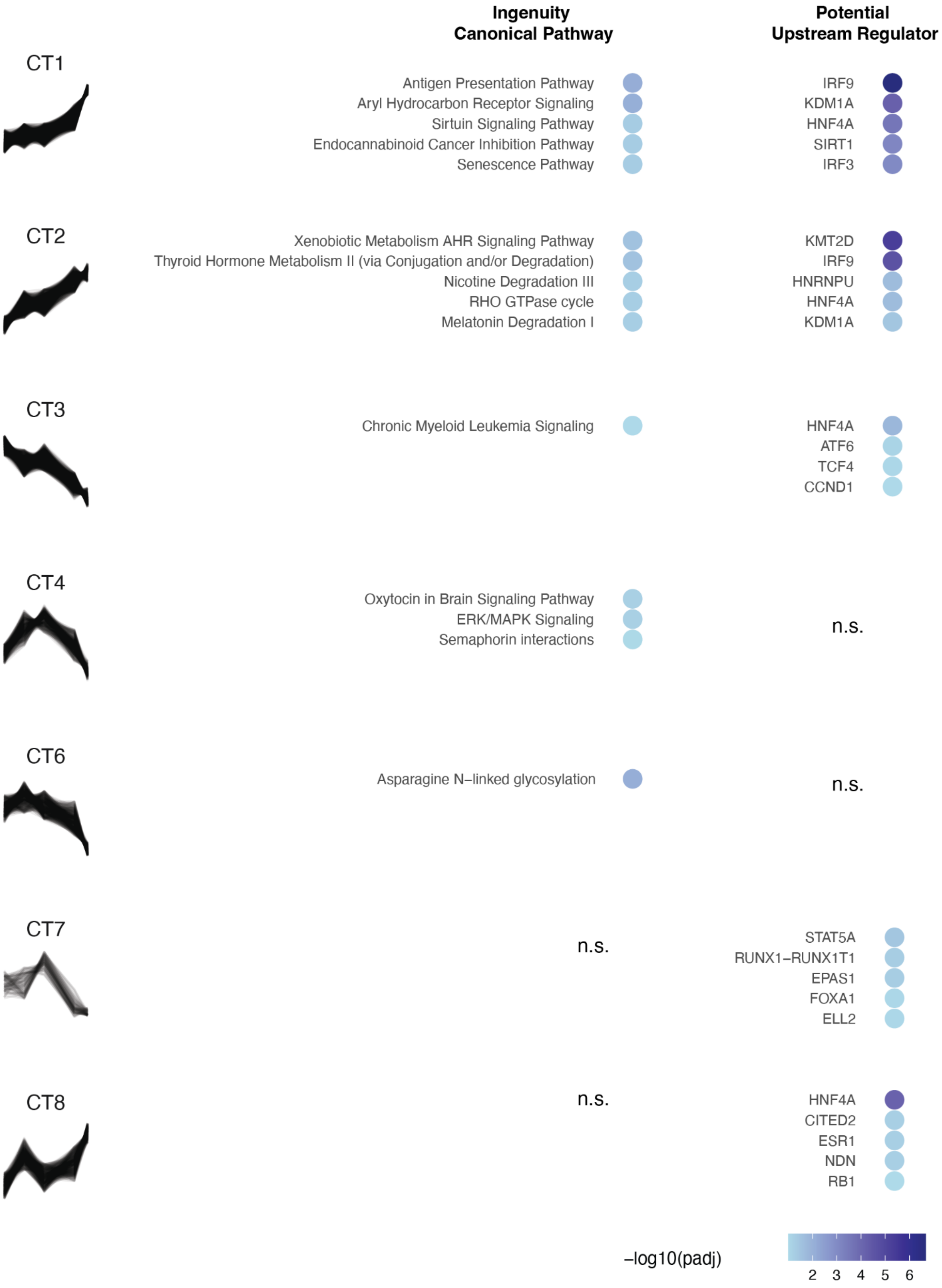


**Supplementary Figure 4.** Top canonical pathways and predicted upstream regulators identified by Ingenuity Pathway Analysis for host genes associated with differentially expressed TEs from different TE clusters. Only results significant after Benjamini-Hochberg correction are shown. n.s., not significant.

a)

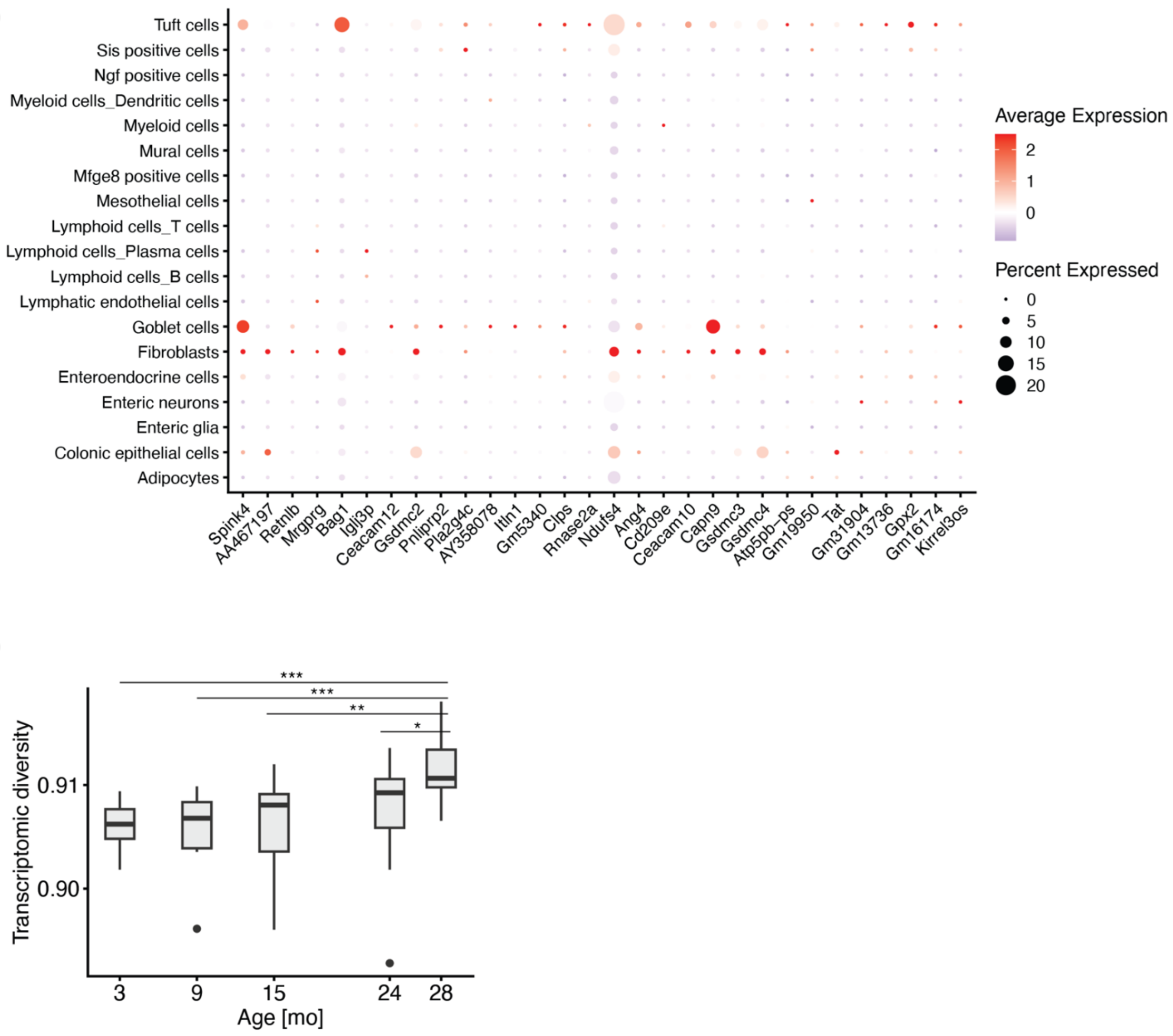


b)



**Supplementary Figure 5.** a) Visualization of the expression patterns of the top 30 genes negatively correlated with transcriptomic diversity in single-cell RNA-seq data generated by Zhang *et al.* (1). Dot size represents the percentage of cells expressing each gene within the indicated group, and dot color represents the scaled average expression level. b) Transcriptomic diversity based on TE expression during aging in the mouse colon.

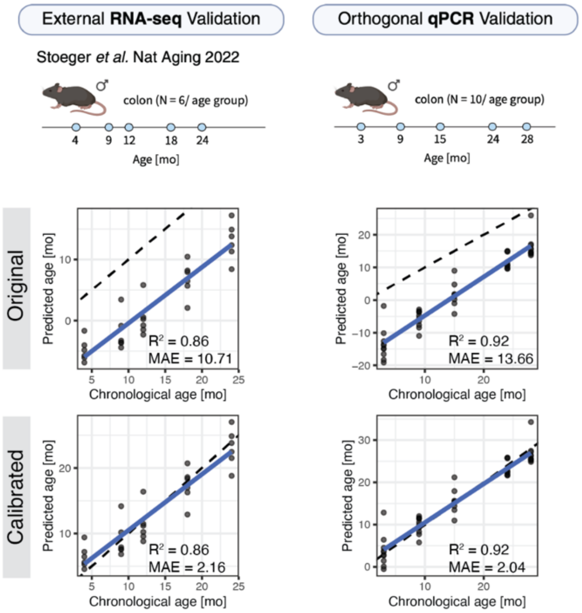


**Supplementary Figure 6.** Datasets used for model validation (top) and predicted age versus chronological age before (middle) and after (bottom) linear recalibration. The upper panel was created with BioRender. $R^2$, squared Pearson correlation between predicted and chronological age; MAE, mean absolute error; dashed line, identity line (y = x); blue line, linear fit.
